# Genomic structural and transcriptional variation of Oryctes rhinoceros nudivirus (OrNV) in Coconut Rhinoceros Beetle

**DOI:** 10.1101/2020.05.27.119867

**Authors:** Kayvan Etebari, Rhys Parry, Marie Joy B. Beltran, Michael J. Furlong

## Abstract

Oryctes rhinoceros nudivirus (OrNV) is a large circular double-stranded DNA virus which has been used as a biological control agent to suppress Coconut Rhinoceros Beetle (*Oryctes rhinoceros*) in Southeast Asia and the Pacific Islands. Recently a new wave of *O. rhinoceros* incursions in Oceania in previously non-infested areas is thought to be related to the presence of low virulence isolates of OrNV or virus tolerant haplotypes of beetles. In this study, chronically infected *O. rhinoceros* adults were field collected from the Philippines, Fiji, Papua New Guinea and the Solomon Islands. We extracted total RNA from these samples to investigate the global viral gene expression profiles and comparative genomic analysis of structural variations between the four different populations. Maximum likelihood phylogenic analysis indicated that OrNV strains from the Solomon Islands and the Philippines are closely related to while OrNV strains from PNG and Fiji formed a distinct adjacent clade. We detected several polymorphic sites with a frequency higher than 35% in 892 positions of the viral genome. The highest number of structural variants, including single nucleotide variants (SNV), insertion, deletion and non-synonymous mutations, were found in strains from Fiji and PNG when compared to complete recently sequenced Solomon Islands OrNV reference genome. Non-synonymous mutations were detected in several hypothetical proteins, and 15 nudivirus core genes such as *OrNV_gp034* (DNA Helicase), *lef-8, lef-4* and *vp91*. For examination of the global gene expression profile of OrNV in chronically infected populations, we found limited evidence of variation between geographic populations. Only a few genes such as *OrNV_gp01* (DNA polymerase B), *OrNV_gp022* and *OrNV_gp107* (Pif-3) were differentially expressed among different strains. Additionally, small RNA sequencing from the Solomon Islands population suggests that OrNV is targeted by the host RNA interference (RNAi) response with abundant 21nt small RNAs. Additionally, we identified a highly abundant putative 22 nt miRNA from the 3’ of a pre-miRNA-like hairpin originating from *OrNV-gp-098*. These findings provide valuable resources for future studies to improve our understanding of the OrNV genetic variation. Some of these structural changes are specific to the geographic population and could be related to particular phenotypic characteristics of the strain, such as viral pathogenicity or transmissibility, and this requires further investigation.

## 1. Introduction

Oryctes rhinoceros nudivirus (OrNV) (Genus *Alphanudivirus*) belongs to the diverse *Nudiviridae* family of rod-shaped viruses with large circular double-stranded DNA genomes which have a nuclear replication strategy in arthropod hosts. The typical pathology observed in nudivirus infected insects, and crustaceans suggest lethal infections in larvae and chronic disease in adults of various terrestrial and aquatic species [1, 2]. The genome of OrNV encodes 130-140 genes [3, 4]. The coconut rhinoceros beetle (*Oryctes rhinoceros*), which is considered a globally invasive species [5] is a pest of palm trees and a severe threat to livelihoods in tropical Asia and the Pacific Islands. *Oryctes rhinoceros* is indigenous to South-East Asia, and OrNV was initially discovered and isolated from insects in Malaysia in 1963, providing a biological control agent for of *O. rhinoceros* that has been successfully used across the region for 50 years [6].

New incursions of the pest into previously *O. rhinoceros-free* countries and territories have been reported in recent years, beginning in Guam in 2007 and followed by Hawaii (2013), Solomon Islands (2015) and most recently Vanuatu and New Caledonia in 2019. It has been suggested that this resurgence and spread of the pest is related to OrNV resistant haplotypes, “CRB-G”, which are identified by single nucleotide polymorphisms in the Cytochrome C Oxidase subunit I (*CoxI*) [7]. Since the release of OrNV into the Pacific, there have been some reports that indicate possible failures of the biocontrol agent including, repeated outbreaks of the beetle, the inconsistency of mortality caused by OrNV and lack of disease symptoms or low incidence of the virus in wild populations [8]. Accordingly, the presence of low virulence isolates of virus or tolerate haplotypes of beetles in different geographical regions has been proposed [9, 10]. Several genomic variants of OrNV have been reported across the region based on endonuclease restriction analyses [11, 12]. Still, the actual genetic diversity present within the wild type populations of OrNV remains unknown. Crawford and Zelazny (1990) detected the first viral structural variations four years after the initial release of different isolates of the virus in the Maldives. Using a restriction endonuclease digestion approach, they described some genomic alterations due to insertion and point mutation and also documented recombination between two of the strains of the virus released, suggesting rapid evolution of the virus [11].

Geographical populations of the virus may vary in virulence due to their genetic diversity. It has been reported that the failure of the virus to maintain control of *O. rhinoceros* in Malaysia may be due to low virulence of endemic OrNV strains [12–14]. The successful establishment of OrNV in *O. rhinoceros* populations is also critically dependent on virus transmissibility, and this too is likely to have a genetic basis. Different levels of transmission have been quantified for OrNV across the region. Still, too little information is available to understand the genetic basis of differences between the virulence and transmissibility of different geographical OrNV strains. Releasing several strains of the virus, as an essential component of pest management strategies, potentially intensifies the genetic diversity of OrNV in the region.

Previous studies have found widespread genomic variations in some large DNA viruses like baculoviruses [15, 16]. Many comparative analyses performed on baculovirus-host interaction models suggest that the differences in virulence and transmissibility involve multiple molecular pathways. Nonetheless, some mutations in key genes can be responsible for low transmissibility or virulence in a particular phenotype of the virus [17, 18]. A single wild type nuclear polyhedrosis virus (NPV) infected *Panolis flammea* caterpillar was found to contain 24 genotypic variants based on restriction fragment length polymorphism [19]. However, the fitness cost associated with genetic diversity in the population and whether or not this genetic diversity can be preserved in the population is unclear. Some non-synonymous mutations in core genes might have a significant impact on the virus fitness within different populations [16]. Although structural variants with low frequency can also be maintained within the population regardless of their fitness cost and can be a dominant genotype when suitable conditions arise [16].

The successful infection of the host gut tissue by OrNV particles is a crucial step that determines the success of the viral infection in *O. rhinoceros*. Many host and viral genes are involved in this critical event, as the midgut represents the first line of cellular defence against viral infection. A complex of proteins called *per os* infectivity factors (PIFs) mediate the binding and entry of virus particles into midgut epithelial cells. Undoubtedly, successful nudivirus infection involves a very complicated and coordinated expression of all viral genes. Any transcriptional variation or non-synonymous mutations of viral genes which disable the normal function of these proteins could reduce virus virulence and interrupt its vertical or horizontal transmission.

Unfortunately, lack of information on the global gene expression pattern of OrNV in persistently (or chronically) infected insect hosts or even in cultured insect cells prevents us from having a clear view of host-pathogen interactions at the viral gene transcriptional level. We recently detected an extremely high prevalence of viral infection in *O. rhinoceros* adult specimens across the region, even in those host haplotypes (CRB-G) which believed to be resistance to OrNV (unpublished data). Here we present the first genome-wide study of the genomic diversity of OrNV strains across the Pacific. This study generated a compendium of OrNV structural variants and gene expression profiles in different geographical strains which infect the different mitochondrial lineages (CRB-G, CRB-S and CRB-PNG) of *O. rhinoceros*. Furthermore, some of these structural changes are specific to different geographic populations and could be related to particular phenotypic characteristics of the strain, such as viral pathogenicity or transmissibility.

## 2. Methods

### 2.1 Sample collection and processing

Adult female Coconut Rhinoceros Beetles (*O. rhinoceros*) were collected from the Philippines (Los Baños) and three different South Pacific countries, Fiji (Sigatoka, Viti Levu), Papua New Guinea (Kimbe, New Britain) and Solomon Islands (Honiara, Guadalcanal) using traps baited with aggregation pheromone (Oryctalure, P046-Lure, ChemTica Internacional, S. A., Heredia Costa Rica) between June-October 2019.

Individual female insects were disinfected by soaking in 75% Ethanol for 30s and then rinsing in phosphate-buffered saline (PBS) before their gut tissues were dissected, removed and preserved in an RNA stabilisation reagent (RNAlater^®^, QIAGEN). For each individual, some fat body and small pieces of gut tissues were also preserved separately in 95% ethanol. Preserved specimens were shipped to the University of Queensland, Brisbane, Australia for further analysis. The preserved gut tissues were kept at −80 °C upon arrival. For examination of virus infection status and mitochondrial haplotype, DNA was extracted from fat bodies and gut fragments using Qiagen Blood and Tissue DNA extraction kit according to manufacturers instruction.

Presence of the OrNV was confirmed by successful amplification of a 945 bp product using the OrV15 primers that target *OrNV-gp083* gene [20] and by Sanger sequencing of each PCR product. For confirmation of the mitochondrial haplotype, a small fragment of *CoxI* gene was amplified in these individuals with the primers LCO1490 and HCO2198 [21]. PCR products were sequenced bi-directionally using an ABI3730 Genetic Analyser (Applied Biosystems) at Macrogen Inc. Seoul, South Korea.

The gut from OrNV positive individuals was further dissected and transferred to Qiazol lysis reagent for RNA extraction according to manufacturer’s instructions (QIAGEN; Cat No.: 79306). The RNA samples were treated with DNase I for 1hr at 37°C and then their concentrations were measured using a spectrophotometer and integrity was ensured through analysis of RNA on a 1% (w/v) agarose gel. After checking the RNA quality, total RNA samples were submitted to the Novogene Genomics Singapore Pte Ltd for RNA sequencing (RNA-seq). The PCR-based cDNA libraries were prepared using NEBNext^®^ Ultra™ RNA Library Prep Kit and were sequenced using Novaseq6000 (PE150) technology with the insert size between 250-300 bp.

### 2.2 Bioinformatic analysis

The CLC Genomics Workbench version 12.0.1 was used for bioinformatics analyses. All libraries were trimmed from any vector or adapter sequences remaining. Low quality reads (quality score below 0.05) and reads with more than two ambiguous nucleotides were discarded. For identification of the virus consensus from each sample, clean reads mapped to a recently reported Oryctes rhinoceros nudivirus complete genome sequence (MN623374) [4]. We implemented strict mapping criteria (mismatch, insertion and deletion costs: 2: 3: 3 respectively). The minimum similarity and length fraction of 0.9 between a mapped segment, and the reference was allowed in mapping criteria. We also permitted minimum 5 reads as low coverage threshold and inserted N as ambiguity symbols for handling the low coverage region. The quality score for each nucleotide has been used to resolve any conflict. For production of multiple sequence alignments, we used MUltiple Sequence Comparison by Log-Expectation (MUSCLE v3.8.425) to compare OrNV whole-genome consensus sequence from different libraries.

#### 2.2.1 Structural variant and phylogenetic analysis

To detect structural modifications among different OrNV infected *O. rhinoceros* populations, CLC Genomic Workbench low-frequency variant detection tool, which is based on a neighbourhood quality standard (NQS) algorithm was used. Variants such as single nucleotide variant (SNV) multiple nucleotide variant (MNV), insertion, deletion and amino acid modification were then called based on neighbourhood radius (5), a maximum number of gaps and mismatches (2), the minimum average quality of surrounding bases (15), and minimum quality of the central base (20). We also used a minimum coverage filter of 10 and a minimum variant frequency of 35% to exclude sequencing artefacts. The final alignments for downstream analysis were OrNV 11×126,183nt and for the partial *CoxI* 11×621nt. For the construction of the maximum-likelihood phylogenetic tree of both host mitochondrial DNA and OrNV, we selected a nucleotide substitution model based on the consensus outcome of hierarchical likelihood ratio test (hLRT), Bayesian information criterion (BIC), Akaike information criterion (AIC). For the OrNV phylogenetic inference the General Time Reversible (GTR) +G rate variation (4 categories), +T topology variation was selected from all tests. For the mitochondrial CoxI the Hasegawa-Kishino-Yano (HKY) substitution model with +T topology variation was selected. The maximum-likelihood phylogeny was then constructed using CLC Genomics Workbench with a Neighbour Joining starting tree construction method under the nucleotide-substitution models mentioned above with 1000 bootstraps. Resultant trees were exported and visualised using FigTree version 1.4 (A. Rambaut; http://tree.bio.ed.ac.uk/software/figtree/).

#### 2.2.2 Viral gene expression profiling

For examination of OrNV gene expression profiles in different *O. rhinoceros* populations, clean reads were mapped to the OrNV genome using the RNA Sequencing Tool on CLC genomics workbench. Expression values for all OrNV genes were calculated as total mapped read count and then normalised to reads per kilobase of transcript per million mapped viral reads (RPKM). The most highly expressed OrNV genes in *O. rhinoceros* gut tissue in different populations were identified and ranked. The viral gene expression profile was created based on calculated RPKM values. For comparison of viral gene expression analysis among different OrNV infected *O. rhinoceros* populations, hierarchical clustering analysis with Euclidean distance metric on the log2-transformed RPKM value (normalised expression value) were performed. Differential expression profiles between the four different populations were modelled using a separate Generalised Linear Model (GLM). The use of the GLM formalism allows the fitting of curves to expression values without assuming that the error on the values is normally distributed. The Wald test was also used to compare each sample against its control group to test whether a given coefficient is non-zero. We considered genes with more than 2-fold changes and a false discovery rate (FDR) of less than 0.05 as significantly modulated viral genes. For further functional annotation of OrNV hypothetical proteins, we search for a conserved domain through IntrProScan database [22]. We also used EggNOG version 4.5 to run pairwise orthology prediction on Cluster of Orthologous Groups [23].

#### 2.2.3 Viral derived small RNA analysis and OrNV miRNA prediction

For analysis of the host RNAi response to OrNV, a small RNA library was generated from a virus-infected individual from the Solomon Islands using the NEBNext^®^ Multiplex Small RNA Library Prep Kit for Illumina^®^ at the Novogene Genomics Singapore Pte Ltd. The purified cDNA libraries were sequenced on Novaseq 6000 (SE50) and raw sequencing reads were obtained using Illuminas Sequencing Control Studio software. Raw data were stripped of adapters and reads with quality score above 0.05, and less than two ambiguous nucleotides were retained. Reads without 3’ adapters and also reads with less than 16 nt were discarded. The clean reads mapped to the recently sequenced OrNV genome (MN623374). For examination of potential viral derived miRNAs of OrNV we used miRDeep v2.0.0.8 [24] hosted on the Mississippi Galaxy instance (https://mississippi.sorbonne-universite.fr/). We screened the genome consensus of all OrNV geographical strains for potential miRNA binding sites with miRanda algorithm [25]. This tool considers matching along the entire miRNA sequence, but we ran the program in strict mode, which demands strict 5’ seed pairing.

## 3. Results and Discussion

### 3.1 Evidence of co-evolution of OrNV strains with their *O. rhinoceros* hosts

In this study *O. rhinoceros* adults were collected from the Philippines in their native region and from Fiji (Sigatoka, Viti Levu), Papua New Guinea (Kimbe, New Britain) and the recently invaded Solomon Islands (Honiara, Guadalcanal) in non-native regions in the South Pacific Islands (Figure 1A). The total RNA from adults *O. rhinoceros* infected with wild type OrNV were sequenced, and in total, 42,678,016 Illumina paired-end reads mapped to the OrNV genome. The percentage of reads mapped to the virus genome in each library was between 0.53-18.92% of the total reads (Table 1). We used the recently sequenced Solomon Islands strain of OrNV genome (MN623374) as a reference in this study [4], and the viral genome consensus sequences have been generated for each library (Figure S1).

**Figure 1:**
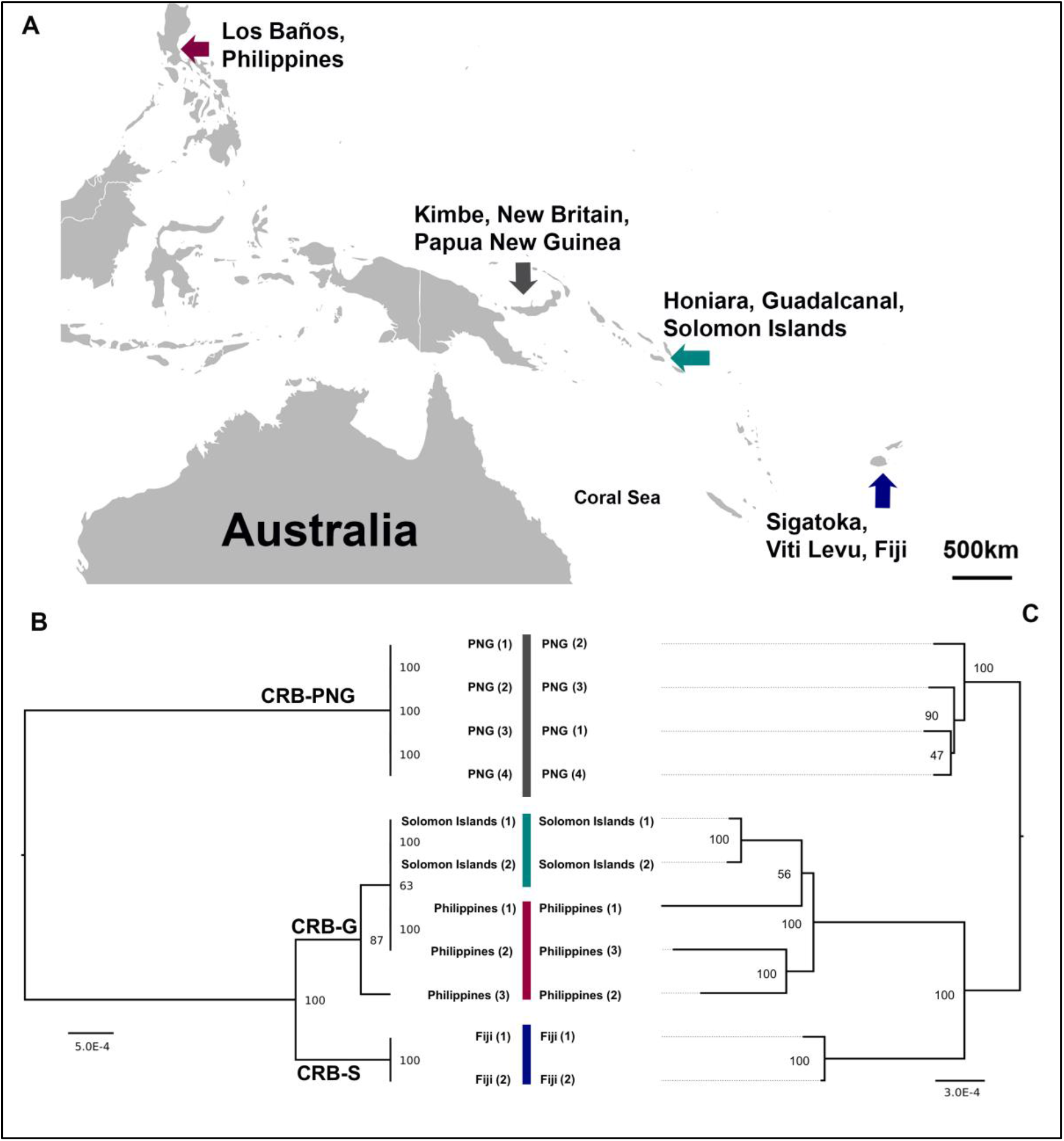
Geographic locations of Coconut Rhinoceros beetles (*O. rhinoceros*) samples chronically infected with OrNV from Southeast Asia and the pacific islands **(A)**. Host mitochondrial maximum likelihood phylogeny based on *O. rhinoceros CoxI* gene sequence. Four different haplotypes of *O. rhinoceros*, in three major haplotype clades (CRB-PNG, CRB-S and CRB-G) were identified **(B)** Maximum likelihood phylogeny analysis of OrNV strains **(C)**. Samples from Solomon Islands and Philippines are closely related to each other while individuals from PNG and Fiji are separated from them and located in another clade. Branch length indicates nucleotide substitutions per site, trees bootstrapped 1000 times. Host phylogenetic tree is midpoint rooted and

**Table 1.** The distribution of viral reads among different RNAseq libraries

| Population | Sample ID | Reads mapped to viral genome | Viral reads in library (%) | Gene (%) | Intergenic (%) |
| --- | --- | --- | --- | --- | --- |
| **Fiji** | F5 | 465,294 | 0.91 | 94.58 | 5.42 |
|  | F6 | 2,498,061 | 6.34 | 93.63 | 6.37 |
| **Philippines** | PH2 | 8,188,367 | 18.92 | 94.08 | 5.92 |
|  | PH4 | 6,129,400 | 13.22 | 93.69 | 6.31 |
|  | PH8 | 525,590 | 0.93 | 92.88 | 7.12 |
| **PNG** | PNG2 | 1,361,190 | 3.21 | 93.31 | 6.69 |
|  | PNG4 | 5,421,019 | 12.65 | 93.58 | 6.42 |
|  | PNG5 | 4,919,054 | 11.87 | 94.1 | 5.9 |
|  | PNG6 | 5,516,617 | 13.62 | 93.28 | 6.72 |
| **Solomon Islands** | SI24 | 7,406,766 | 14.33 | 94.04 | 5.96 |
|  | SI30 | 246,658 | 0.54 | 91.54 | 8.46 |

Maximum likelihood phylogeny analysis of four different wild type populations of OrNV based on their complete genome sequence consensus generated two major clades. These data show that samples from Solomon Islands and Philippines are closely related to each other while samples from PNG and Fiji are separated from them and located in another clade (Figure 1). The phylogenic analysis of field-collected *O. rhinoceros* based on the host partial *CoxI* gene revealed three major mitochondrial lineages, CRB-G, CRB-S and CRB-PNG. All *O. rhinoceros* individuals collected from PNG (Kimbe, New Britain) and Fiji belonged to the mitochondrial lineages CRB-PNG and CRB-S, respectively. The individuals from the Philippines and the recently invaded Solomon Islands belonged to the CRB-G clade (Figure 1C). The virus strains from the Solomon Islands and the Philippines also belonged to one major clade (Figure 1B). Harmonious topologies of both host and OrNV strains provide evidence of ongoing co-evolution of OrNV strains with their *O. rhinoceros* hosts.

Previously Marshall *et. al*., (2017) reported that the natural infection rate of OrNV ranges from 45.2% to 64.5% in *O. rhinoceros* from a wide range of countries in Southeast Asia and Pacific. However, they couldn’t detect the presence of the OrNV in specimens from the Solomon Islands, Philippines and Oahu which were believed to be the CRB-G haplotype. The source of the virus found in our 2019 collection from the Solomon Islands is not known [4]; it could have been introduced deliberately as a biological control agent or along with an incursion of infected beetles from neighbouring islands (CRB-PNG from PNG). However, due to high the high level of similarity and lack of any evidence that OrNV (Philippines strain) was introduced into the country as a biological control agent, our work suggests that the OrNV came to the Solomon Islands with an incursion of infected CRB-G that originated in Southeast Asia.

### 3.2 Whole genome structural variant analysis of OrNV strains

We detected several SNPs with a frequency higher than 35% in 892 positions of the 125917 bp viral genome sequence, which found in pair-end reads positioned in both orientations of the genome. In samples collected from the Solomon Islands, 87-141 variants were identified in which deletion is their major variation when compared to the newly sequenced genome reference (MN623374). The maximum number of structural variants such as single nucleotide variants (SNV), insertion, deletion and the number of variants in amino acid (non-synonymous mutations) were found in individuals from Fiji and PNG (Figure 2). We detected 12-17 proteins with amino acid modifications in samples collected from the Solomon Islands while this number reached to 61-63 proteins in samples from Fiji and PNG. OrNV in the Philippines showed more similarity to OrNV in individuals from Solomon Islands based on the number of identified SNV and non-synonymous modifications. Some of the structural variants we identified could be involved in differences in viral transmission and virulence.

**Figure 2:**
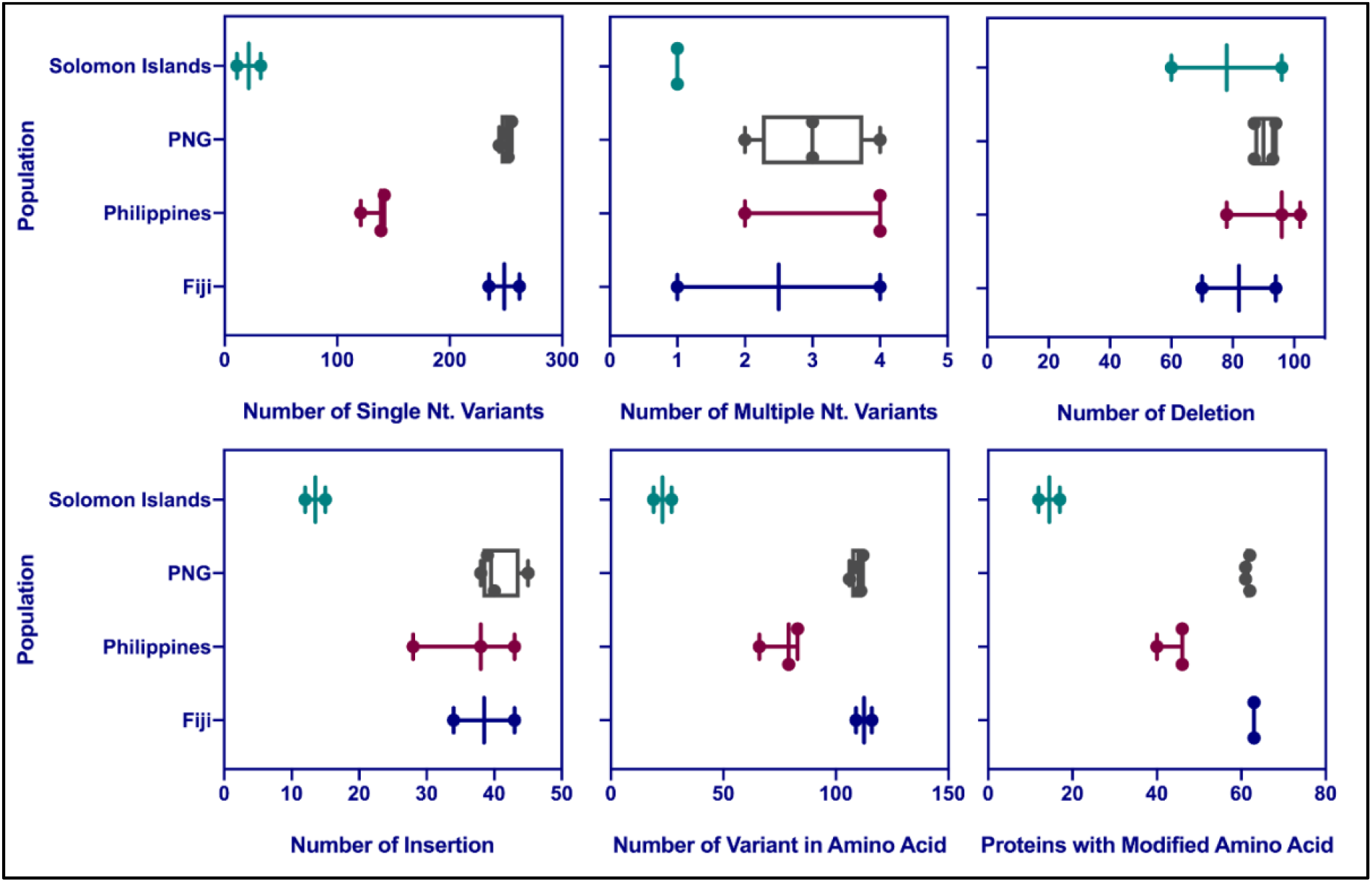
Genomic structural variants of different geographical populations of OrNV when compared to the newly sequenced reference genome of a Solomon Islands strain.

Structural variants have been found in both coding and non-coding regions of the OrNV genome. In all individuals from Fiji, Philippines and PNG, the number of variants in coding regions is higher than the number in the non-coding part of their genomes (Figure 3). Generally, mutations in coding regions are likely to have a significant impact on the genome, whereas, in non-coding regions, such mutations are more likely to be neutral. However, certain non-coding regions of baculoviruses could be important components of the viral genomes. [17].

**Figure 3:**
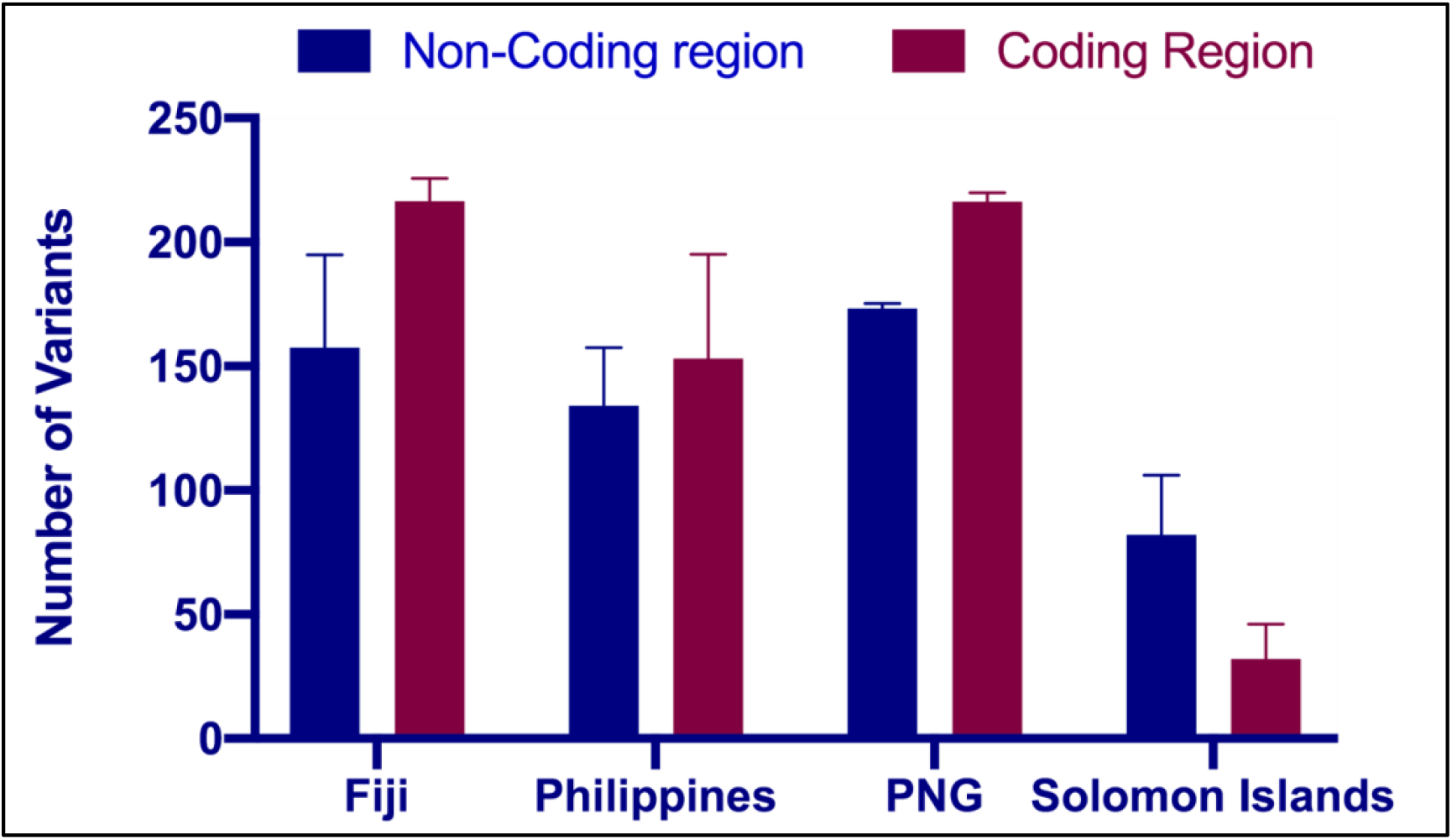
The number structural variants in coding and non-coding regions of different strains of OrNV.

Non-synonymous mutations which do result in amino acid modification and could potentially interrupt protein function were detected in 15 nudivirus core genes and several hypothetical proteins (Table 2 and Table S1). We found up to four amino acid modifications in DNA Helicase (QHG11272.1) among individuals from different populations. A non-synonymous mutation (c.5C>T) in *OrNV_gp034* caused an amino acid modification in DNA helicase in and individual sample from Solomon Islands. This amino acid substitution from Threonine to Methionine can also change the biochemical properties (polarity changed from neutral polar to non-polar) of this critical protein, and therefore it might affect protein function and possibly virus performance. Other non-synonymous mutations in this gene do not change the polarity of the protein. One of these amino acid changes (p.Leu211Phe) is specific to samples isolated from Fiji. Previous studies in baculoviruses showed that this gene is associated in host range expansion. Mutations in the genes involved in DNA replication might affect the rate of virion production and thus might alter the spread of the infection within the host [17]. DNA Helicase is one of the most rapidly evolving genes in Drosophila innubila nudivirus (DiNV) and Kallithea virus and may be important for adaptation to a new host system [26, 27].

**Table 2.** Number of amino acid modifications in OrNV core gene products.

| Gene ID | Gene | Gene Product | Protein NCBI ID | FIJI 1 | FIJI 2 | PHI 1 | PHI 2 | PHI 3 | PNG 1 | PNG 2 | PNG 3 | PNG 4 | SI 1 | SI 2 |
| --- | --- | --- | --- | --- | --- | --- | --- | --- | --- | --- | --- | --- | --- | --- |
| **DNA replication:** |  |  |  |  |  |  |  |  |  |  |  |  |  |  |
| OrNV_gp001 | *dnapol* | DNA polymerase B | QHG11240.1 | 2 | 2 | 4 | 2 | 2 | 2 | 2 | 2 | 2 | 2 | - |
| OrNV_gp034 | *helicase* | DNA helicase | QHG11272.1 | 4 | 4 | 3 | 1 | 2 | 3 | 3 | 3 | 3 | - | - |
| OrNV_gp108 | *helicase* | DNA helicase 2 | QHG11340.1 | - | - | - | 1 | 1 | - | - | - | - | - | - |
| **Transcription:** |  |  |  |  |  |  |  |  |  |  |  |  |  |  |
| OrNV_gp020 | *p47* | p47 protein | QHG11259.1 | - | - | - | - | - | - | - | - | - | - | - |
| OrNV_gp064 | *lef-8* | late expression factor 8 | QHG11301.1 | - | - | 1 | - | - | 1 | 1 | - | - | - | - |
| OrNV_gp096 | *lef-9* | late expression factor 9 | QHG11328.1 | 2 | 2 | 1 | 1 | 1 | 2 | 2 | 2 | 2 | - | - |
| OrNV_gp042 | *lef-4* | late expression factor 4 | QHG11280.1 | 1 | 1 | - | - | - | - | - | - | - | - | - |
| OrNV_gp052 | *lef-5* | late expression factor 5 | QHG11289.1 | - | - | - | - | - | - | - | - | - | - | - |
| OrNV_gp059 | *lef-3* | late expression factor 3 | QHG11296.1 | - | - | - | - | - | - | - | - | - | - | - |
| **Envelope component:** |  |  |  |  |  |  |  |  |  |  |  |  |  |  |
| OrNV_gp017 | *pif-2* | per os infectivity 2 | QHG11256.1 | - | - | - | - | 1 | - | - | - | - | - | - |
| OrNV_gp060 | *pif-1* | per os infectivity 1 | QHG11297.1 | - | - | 2 | - | - | 1 | 1 | 1 | 1 | - | - |
| OrNV_gp107 | *pif-3* | per os infectivity 3 | QHG11339.1 | - | - | 1 | - | - | - | - | - | - | - | - |
| OrNV_gp033 | *pif-4* | 19 kDa protein | QHG11271.1 | - | - | - | - | - | - | - | - | - | - | - |
| OrNV_gp115 | *pif-5* | odv-e56 protein | QHG11347.1 | 1 | - | - | - | - | - | - | - | - | - | - |
| OrNV_gp126 | *p74* | p74 protein | QHG11358.1 | 3 | 3 | 1 | 1 | 2 | 1 | 1 | 1 | 1 | 1 | 1 |
| OrNV_gp004 | *ac81* | Ac81-like protein | QHG11243.1 | - | - | 1 | - | 1 | - | - | - | - | - | - |
| OrNV_gp113 | *p33* | Ac92-like protein | QHG11345.1 | - | - | - | - | - | - | - | - | - | - | - |
| **Capsid component:** |  |  |  |  |  |  |  |  |  |  |  |  |  |  |
| OrNV_gp087 | *38K* | 38K protein | QHG11320.1 | - | - | - | - | - | - | - | - | - | - | - |
| OrNV_gp072 | *ac68* | Ac68-like protein | QHG11307.1 | - | - | - | - | - | - | - | - | - | - | - |
| OrNV_gp030 | *vlf-1* | very late expression factor 1 | QHG11269.1 | 2 | 2 | 2 | 2 | 1 | 2 | 2 | 2 | 2 | - | - |
| OrNV_gp022 | *gp72* | GrBNV_gp72-like protein | QHG11261.1 | 1 | 1 | - | - | - | 1 | 1 | 1 | 1 | - | - |
| OrNV_gp015 | *vp39* | Viral capsid associated protein 39 | QHG11254.1 | - | - | - | 1 | - | - | - | - | - | - | - |
| OrNV_gp106 | *vp91* | Viral capsid associated protein 91 | QHG11338.1 | 2 | 2 | - | 1 | - | 1 | 1 | 1 | 1 | - | - |

We identified non-synonymous mutations in several genes involved in transcription, including three subunits of the RNA polymerase (*lef-4, lef-8* and *lef-9*) (Table 2 and Table S1). Interestingly, a similar event has been reported from AcMNPV [16] and *Spodoptera exigua* multiple nucleopolyhedrovirus (SeMNPV) [17]. Here, amino acid variants in *lef-9* do not change protein polarity. Still, non-synonymous mutations in *lef-8* and *lef-4* had an impact on biochemical properties of the protein in some individuals from Fiji and PNG. *OrNV_gp64* which encode lef-8 protein is also overexpressed in OrNV collected from Fiji (Fold change 4.38, FDR: 0.048) and the lowest expression value was detected in individuals from the Philippines. *lef-4* is an RNA capping enzyme and was originally identified as being essential for late transcription. A deletion mutant in the BmNPV *lef-8* homolog (Bm39) was not viable, and inactivation of BmNPV *lef-8* prevented virus replication [28, 29]. Theze et al (2014) also suggested that mutations in the subunits of the viral RNA polymerase, *lef-9* and *p47* ORFs are associated with vertical transmission and pathogenicity. We did not detect any mutations in *OrNV_gp020*, which encodes p47 protein, in individuals the sequenced from across the pacific suggesting that this gene is essential for OrNV replication. However, it is challenging to definitively link specific mutations to virus performance because virulence and virus performance is derived from the combination of the genetic material present at the whole-genome scale.

Genes involved in primary infection encoding essential components of the *per os* infectivity complex such as *pif-1, pif-2, pif-3* also have some non-synonymous mutations in their sequences across the different geographical population of OrNV (Table 2 and Table S1). The products of *pif* genes, PIF proteins, which are conserved across the baculoviruses and nudiviruses, bind to insect midgut cells and are essential for infection. We only detected one amino acid modification in *pif-3* and *pif-2* in one of the samples collected from the Philippines. There is a conserved amino acid change (p.Val184Ile) in *pif-1* of all individuals from the PNG population, but it has no impact on protein polarity. Chateigner et al (2015) suggested that the presence of different variants of the PIF proteins in the AcMNPV might allow the binding of these proteins to a wider range of host cell, which could be advantageous for establishment of viral infection.

*OrNV_gp126* which encodes p74 protein is another gene with a role in *per os* infectivity, and we detected amino acid changes in all individuals. This protein is critical in host-pathogen interactions in the insect midgut, and its deletion prevents the recombinant virus transmission in insects, but the mutation does not affect replication in cultured cells [30]. Two of the amino acid changes were specific to the Fiji population and altered protein polarity.

The very late factors (VLF) appear to be a structural, conserved protein that is present in both baculoviruses (BV and ODV) and nudiviruses and which is required for the production of nucleocapsids. Several deletion/insertions of serine (p349^350) and glutamine (p644^645) have been detected in *vlf-1* in all the OrNV samples; this could be due to a sequencing error (homopolymer stretches) or the presence of actual mutation sites (Table 2 and Table S1). We also detected two mutation sites in *OrNV_gp106*, which encodes VP91 protein. One of those mutations (p.Glu564Asp) is specific to OrNV from the Solomon Islands and the Philippines, and another (p.Ala117Thr) is specific to OrNV from Fiji. Changing alanine to threonine also alters protein polarity from non-polar to polar, and it could cause critical conformational changes affecting protein structure. Theze *et al* (2014) showed that some variants of *vp91* in SeMNPV could be associated with pathogenicity [17].

### 3.3 Limited evidence of OrNV gene expression variation between chronically infected populations

The principal component analysis of OrNV samples from different countries based on overall viral gene expression profiles can differentiate each of the OrNV populations. Two samples from Fiji had similar profiles, and their expression pattern is slightly different from the other populations (Figure 4). Two individuals from the Solomon Islands and Philippines are located outside of their population cluster, probably due to different stages of infection, specific gene expression patterns and low viral reads in those libraries (Figure 4 and Table 1). However, we performed library size normalisation using the TMM (trimmed mean of M values) method and library sizes are then used as part of the pre-sample normalisation. To get a better view of gene expression profile in each sample, we ranked the genes based on their RPKM values and filtered the most highly and poorly expressed gene in each library.

**Figure 4:**
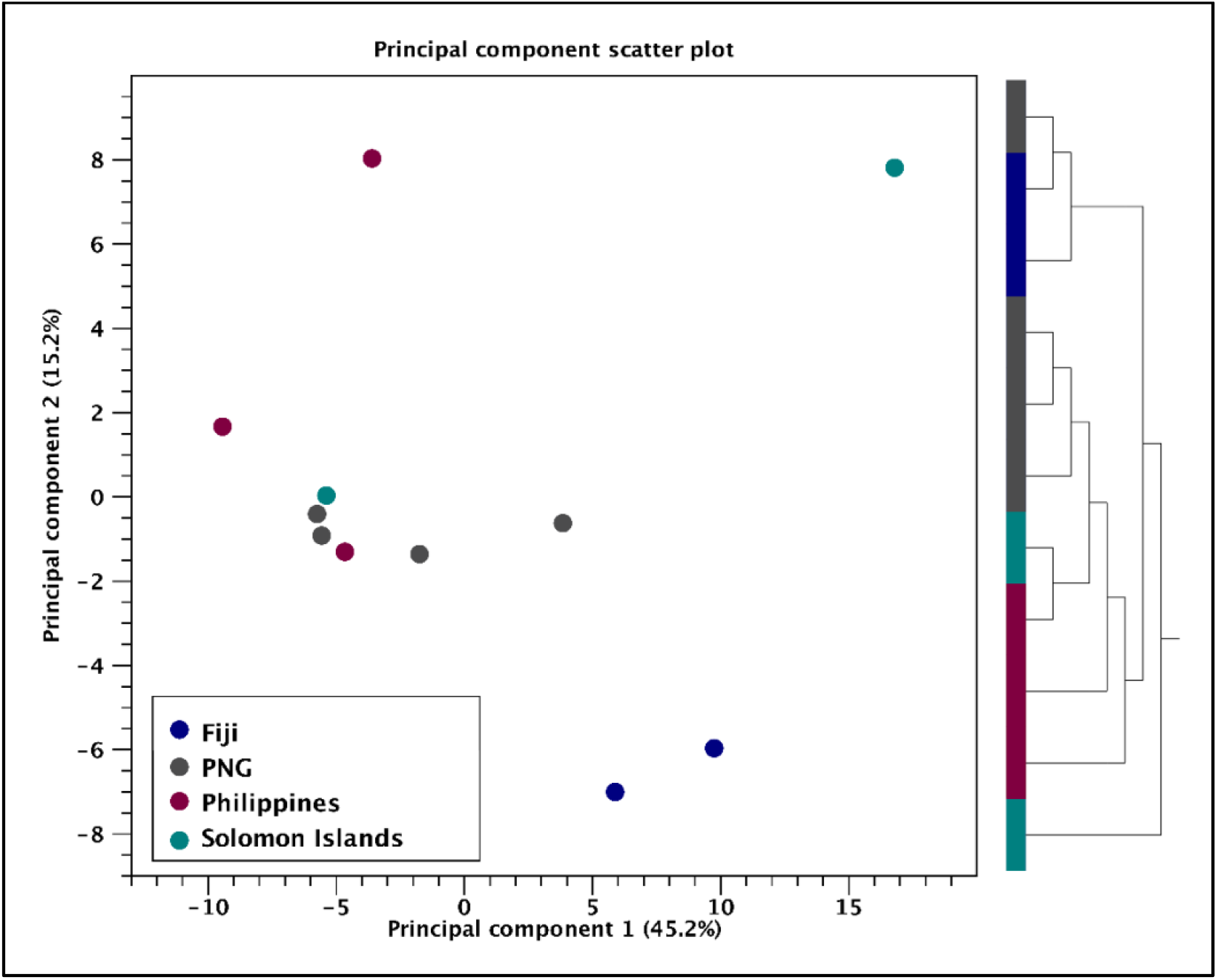
Principal component analysis of OrNV samples from different regions based on the global viral gene expression profile.

*OrNV_gp41* is the most highly expressed gene with an unknown function in all individual libraries. This 342 bp nudivirus specific gene shows significant similarity to GrBNV_gp95 and does not have a known homolog in Baculoviruses. Further sequence analysis using InterProScan and GO database showed that some parts of *OrNV_gp41* gene product are integral components of the hydrophobic region of the membrane (GO:0016021). This virus fragment also integrated into the brown planthopper (*Nilaparvata lugens*) genome [31]. Recently, it has been identified as endogenous nudivirus-like viruses of parasitoid wasps in CcBV (*Cotesia congregate* Bracovirus), CiBV (*Chelonus inanitus* Bracovirus) and its expansion into a gene family in a wasp genome also reported from *Venturia canescens* virus-like particles (VcVLP) [32].

*OrNV_gp022* (GrBNV_gp72-like) is a member of alphanudivirus core genes with unknown function, which is in the top 10 highly expressed genes in all OrNV populations. However, its normalised expression value in the Fiji population is significantly lower than in other samples (Figure 5 and Table S2). Other genes among the top 10 most highly expressed genes are *OrNV_gp015, gp002, gp054, gp012* and *gp023* which encode viral capsid Vp39, a trypsin-like serine protease, glycoprotein gp83-like, odv-e66 and guanylate kinase-like proteins, respectively.

**Figure 5:**
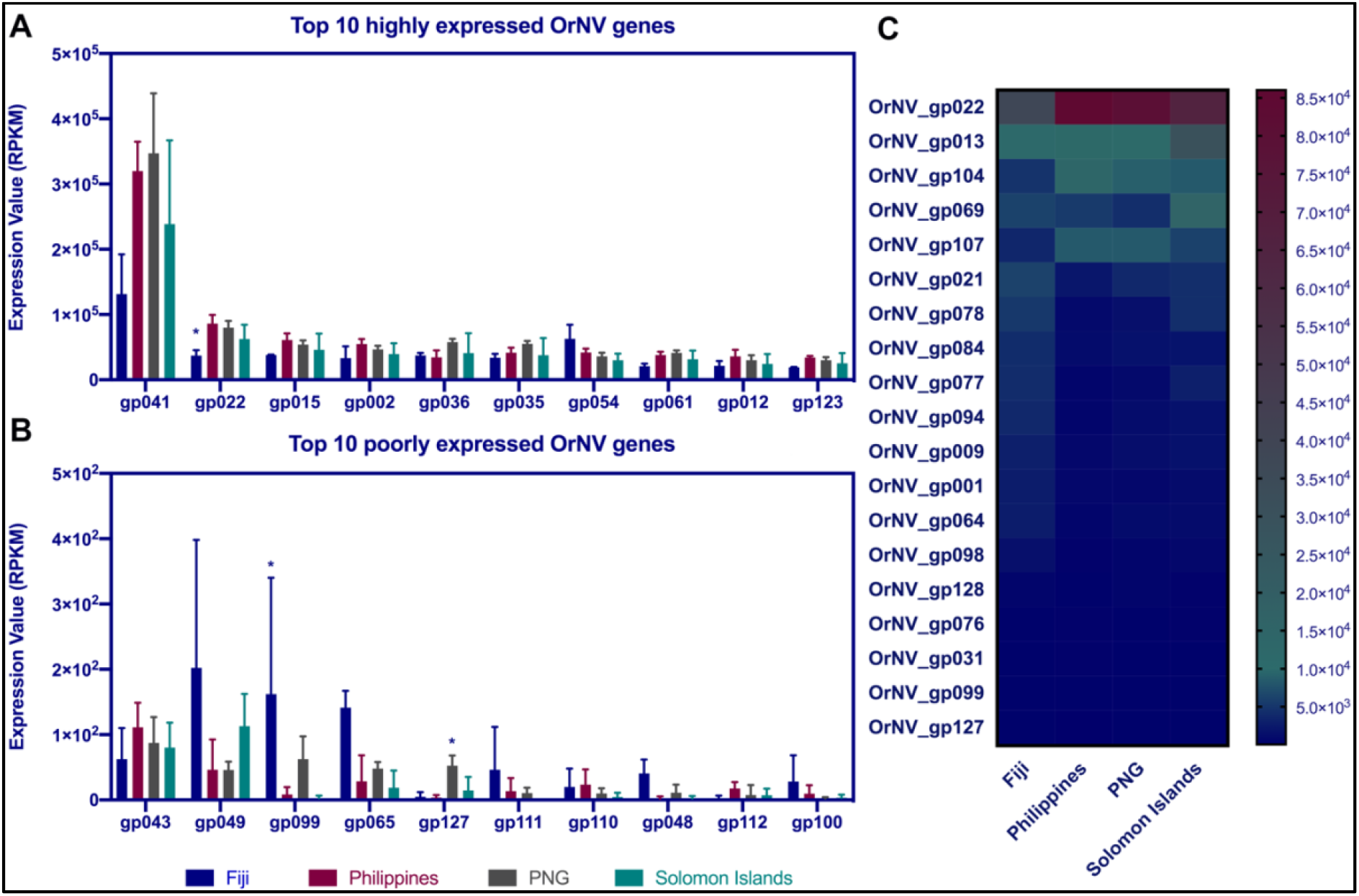
Top 10 highly and poorly expressed OrNV genes in different geographical strains **(A and B)**. Only 19 genes are differentially expressed among four OrNV strains **(C)**.

Trypsin-like serine proteases are non-structural proteins with a conserved domain (cl21584) which are found in several RNA and DNA viruses and cellular organisms [33]. These proteins are essential for virus maturation [34], and they are involved in proteolysis through serine-type endopeptidase activity and post-translational modifications of these enzymes play a crucial role in virus replication [35]. Previously, it has been shown that trypsin-serine proteases have dN/dS above 0.5 between DiNV and OrNV, which indicate that this gene is under purifying selection (dN/dS < 1) and unconstrained evolution or putative adaptation (dN/dS >0.5) [26]. However, we did not detect any amino acid modification in this gene within the different OrNV populations.

Recently, it has been experimentally demonstrated that a nudivirus encoded protein gp83 inhibits the Toll signalling pathway in the fruit fly. Drosophila melanogaster Kallithea virus (KV), which is a new member of family Nudiviridae and DiNV suppress host immune responses by regulating NF-kB transcription factors [36]. This immunosuppressive activity of *gp83* against *D. melanogaster* NF-kB signalling is conserved and probably *gp83* shows similar activity in other members of Nudiviridae. This gene is the second most highly expressed gene in samples collected from Fiji, and its normalised expression value is slightly higher than in other populations (Figure 5).

ODV-E66 protein which has a Chondroitin AC/ alginate lyase (IPR008929) domain, potentially involves viral envelope formation and facilitates viral protein movement during infection by degrading larval peritrophic membrane [37]. This protein has been reported from baculoviruses and other nudiviruses (such as HgNV and Penaeus monodon nudivirus), and it has been suggested that ODV-E66 is a *per os* infectivity factor [2, 38]. Although *odv-e66* is not a top 20 most highly expressed gene in AcMNPV, its expression pattern has a strong negative correlation between *Trichoplusia ni* midgut and *T. ni* cell line (Tnms42). Overexpression in midgut tissue confirms their responsibility for the trafficking of viral proteins during infection [18].

*OrNV_gp107* which encodes pif-3 is highly expressed in OrNV strains from the Philippines, and its expression is 3.56 times higher than in samples collected from Fiji (Figure 5C). Although expression of *pif* genes are necessary for successful infection of gut tissue, it has been shown that over-expression *pif-1* in *Spodoptera frugiperda* NPV is unfavourable for viral population fitness and can cause more mutation in this gene and develop viral genotypes with lack of *pif-1* expressing capabilities [39]. There are several pif genes in OrNV which encode essential components of envelope proteins and contain an N-terminal hydrophobic sequence in combination with several adjacent positively charged amino acids.

Most of the OrNV poorly expressed genes encode hypothetical proteins, and we don’t have enough information about their potential role in the host-pathogen interaction. *OrNV_gp99* which encodes Ac120-like protein is one the most poorly expressed genes in OrNV but their expression was significantly upregulated in samples collected from Fiji and almost undetectable in Philippines and Solomon Islands populations (Figure 5, Table S2). We recently found six nucleotides substitution in *OrNV_gp99* when we compared the Solomon Islands strain with Ma07 from Malaysia, and several amino acid modifications were detected among samples analysed in the current study (Table S1). Homologs of this gene have been found in other baculoviruses and nudiviruses, and it is likely to be nonessential, as an insertion/deletion mutation of this gene in BmNPV (Bm98) had no apparent effect on infectivity [40].

*OrNV_gp01* which encodes DNA polymerase B has low expression value in almost all libraries (Rank 97 out of 130 genes), but it is overexpressed (Fold change 2.7, FDR: 0.002) in samples collected from Fiji (Figure 5C).

### 3.4 OrNV is targeted by the host RNAi response and encodes a putative miRNA

During virus infection in arthropods, two standard small RNA classes; the microRNAs (miRNAs) and small-interfering RNAs (siRNAs) contribute to the antiviral response, promote viral tolerance and in some cases, clearance of virus infections (Reviewed by [41, 42]). During DNA virus infection, the host riboendonuclease III enzyme Dicer-2 cleaves double-stranded RNA (dsRNA) into virus-derived small interfering RNAs (vsiRNAs) 20–22 nt in length [43, 44]. These vsiRNAs are loaded into the RNA-induced silencing complex (RISC), where they target RNA molecules through complementarity, reduce DNA virus gene transcription and ultimately virus replication. The RNAi response in DNA virus infection is subtly different to the RNAi response to RNA virus infections as dsDNA viruses do not require an RNA genome intermediate for replication. As such, the biogenesis of vsiRNAs are abundantly generated from the bi-directional transcription of overlapping gene regions [43] and also secondary RNA structures [45]. Some secondary structures of viral RNA are processed by the canonical host miRNA machinery and appear to be functional miRNAs [46]. These miRNAs influence virus latency in the case of Heliothis zea Nudivirus [47], but can also autoregulate transcription of DNA virus genes [48].

To our knowledge, only one study has examined the virus RNAi response in beetles with virally derived siRNAs demonstrated in the red flour beetle (*Tribolium castaneum*) infected with the single-stranded positive-sense RNA Tribolium castaneum iflavirus (*Iflavirdae*) [49]. To examine the repertoire of small RNA responses to OrNV, the small RNA fraction (16-32nt) of a persistently infected sample from the Solomon Islands was subject to small RNA sequencing and mapped against the OrNV genome. We examined both the size distribution of the viral derived RNA fragments as well as “hot-spot” genomic locations. After quality and adapter trimming of the small RNA sequencing library (Figure 6A) 563,774 of the 11,121,239 small RNA 16-32nt reads (5.07%) could be mapped to both orientations of the OrNV genome. Examination of the size profile of these mapped small RNA reads (Figure 6B) suggested that the majority (57.7%) correspond to the 21nt prototypical Dicer-2 vsiRNAs. After examination of the strand origin of the viral derived small RNA read population, reads originated from both the forward (52.7%) and reverse orientation (47.3%) of the genome more or less evenly. The genomic regions responsible for the high generation of vsRNAs in the OrNV genome suggested that the biogenesis of vsRNAs from the OrNV were unevenly targeted throughout the OrNV genome with “hot spot” regions as previously demonstrated in other DNA-insect RNAi responses [43, 44]. This is evidenced by coverage on both forward and reverse sequences similar to a pattern of “mirroring” of vsiRNA between genome position (Figure 6C).

**Figure 6:**
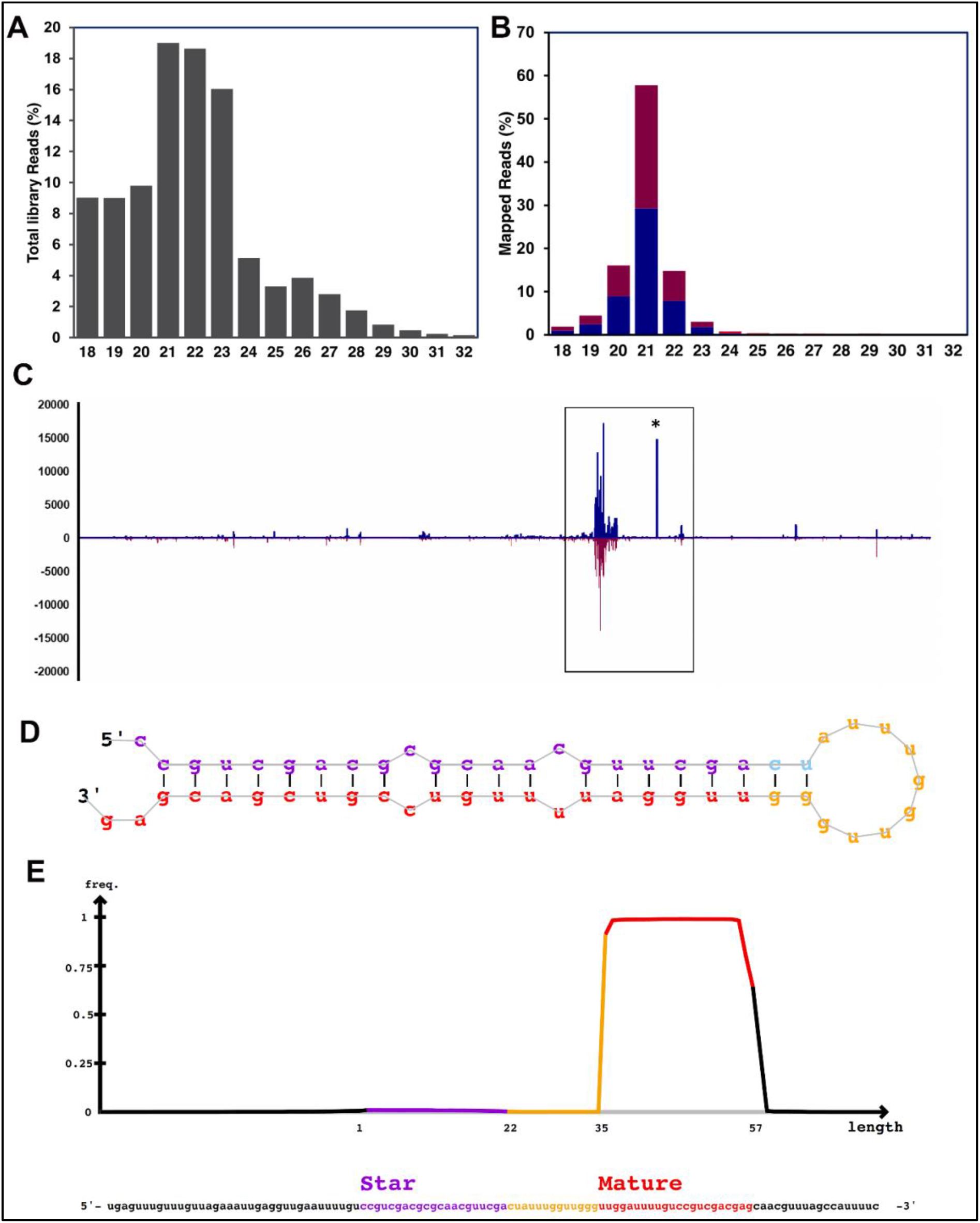
OrNV is targeted by the host RNAi response and encodes a putative miRNA. The length distribution of the small RNA library **(A)**. The length distribution of reads mapped to the OrNV genome which demonstrated a peak at 21nt **(B)** the red and blue bars represent reads mapped to negative and positive strands respectively. Noticeable hot spots were detected in several positions on the OrNV genome **(C)**. The potential miRNA location is shown by an asterisk. OrNV-miR-1 hairpin loop structure **(D)**. The small RNA read density mapped to different parts of potential OrNV precursor miRNA. Significant number of reads mapped to mature OrNV-miR-1-3p **(E)**.

One portion of high coverage with vsRNA reads without a corresponding region of high coverage originating from overlapping negative genome orientation at approximately 85.5kb of the OrNV genome (Figure 6C). As a previous report from *Drosophila* infected with Kallithea Virus (*Nudivirdae*), a close relative of OrNV, suggests that Kallithea Virus encodes a miRNA [50] we examined the possibility that OrNV also potentially produces a miRNA. For this small RNA reads originating from the OrNV genome were analysed by the miRNA prediction tool mirDeep2 [24]. A single putative 22 nt miRNA (UUGGAUUUUGU-CCGUCGACGAG) from the 3’ of a pre-miRNA-like hairpin originating from the genomic location corresponding to a high peak of small RNAs (85464-85520 on *OrNV-gp-098*) (Figure 6C-D). The mature sequence and isomir variants of this putative miRNA (herein described as OrNV-miR-1-3p) were highly abundant in the whole sRNA library and in total considering 2 nucleotide upstream and 5 nucleotide downstream amounted to almost 2.6% of all OrNV reads with (14,595 reads).

Additionally, the mature OrNV-miRNA-1 was the single most abundant sRNA sequence of the OrNV derived small RNAs with the exact 22nt accounting for 4832 reads (0.8%). There were only 65 reads in total associated with −5p RNA “star” sequence (OrNV-miR-1-5p: CCGUCGACGCGCAACGUUCGA) (Figure 6E).

Upon further examination of the miRNA, we found that there was no apparent homology to other coleopteran or arthropod miRNAs as per similarity analysis to miRNAs annotated on miRBase (Release 22.1: October 2018) [51] and the seed region displays no obvious similarity to known miRNAs. Unfortunately, as the genome of *O. rhinoceros* is not currently available, we are not able to conduct a preliminary analysis to potential OrNV-miR-1/host gene mRNA interactions however miRNA could alternatively regulate virus gene expression of OrNV itself [46]. We identified several potential binding sites on the genome of different geographical strains of OrNV (Supplementary Figure S2). Although 49 potential binding sites are universal among all strains, there are some binding sites exclusively identified for Fiji, PNG and the Philippines, which are not detected in other populations (Supplementary Figure S2). While numbers of OrNV-miR-1 are far lower than the 22 nt miRNA reported from Kallithea Virus, which represented >35% of all reads [50], given the evidence that other large dsDNA viruses with nuclear replication strategies also encode miRNAs and the absence of overlapping transcription or high −5p or “star” reads indicates that OrNV-miR-1 is a promising candidate for future laboratory studies in the contribution to long term chronic OrNV persistence in infected *O. rhinoceros* beetles.

## 4. Conclusion

We identified a high number of polymorphic sites among several geographical strains of OrNV, but potentially only a few of these variations in the genome are involved in functional changes. We found some structural variations in OrNV core genes such as *OrNV_gp034* (DNA Helicase), *lef-8*, *lef-4* and *vp91*, which can potentially alter their typical function. These findings provide valuable resources for future studies to improve our understanding of the OrNV genetic variation in different geographic regions and their potential link to virus pathogenicity. However, further genomic analyses of OrNV strains in combination with the characterisation of its effects on the host across multiple populations is necessary to comprehend if such structural or transcriptional variation is responsible for different levels of effectiveness of OrNV as a biological control agent in different countries.

## Data Availability

Deep sequencing data have been deposited in NCBI’s Gene Expression Omnibus and are accessible through GEO series accession numbers GSExxxx and GSExxxx (pending accession numbers).

## Author Contribution

Kayvan Etebari: Conceptualisation, Methodology, Investigation, Formal analysis, Data curation, Visualisation, Writing-Original draft preparation, Funding acquisition

Rhys Parry: Data curation, Formal analysis, Visualisation, Writing-Original draft preparation Marie Joy B. Beltran: Resources

Michael J. Furlong: Conceptualisation, Writing-Reviewing and Editing, Funding acquisition, Project administration, Supervision

## Acknowledgements

This project was supported by the Australian Centre for International Agricultural Research funding (HORT/2016/185) and the University of Queensland (UQECR2057321). We would also like to thank Apenisa Sailo (Ministry of Agriculture, Fiji), Helen Tsatsia (Ministry of Agriculture and Livestock, Solomon Islands) and colleagues from Papua New Guinea Oil Palm Research Association, Dami research station for their assistance in collecting and providing the insect specimens.

## Supplementary Files

**Table S1:** List of all identified structural variants on coding regions of OrNV among all different geographical strains.

**Table S2:** Differentially expressed viral genes among samples collected from the different populations.

**Figure S1:**
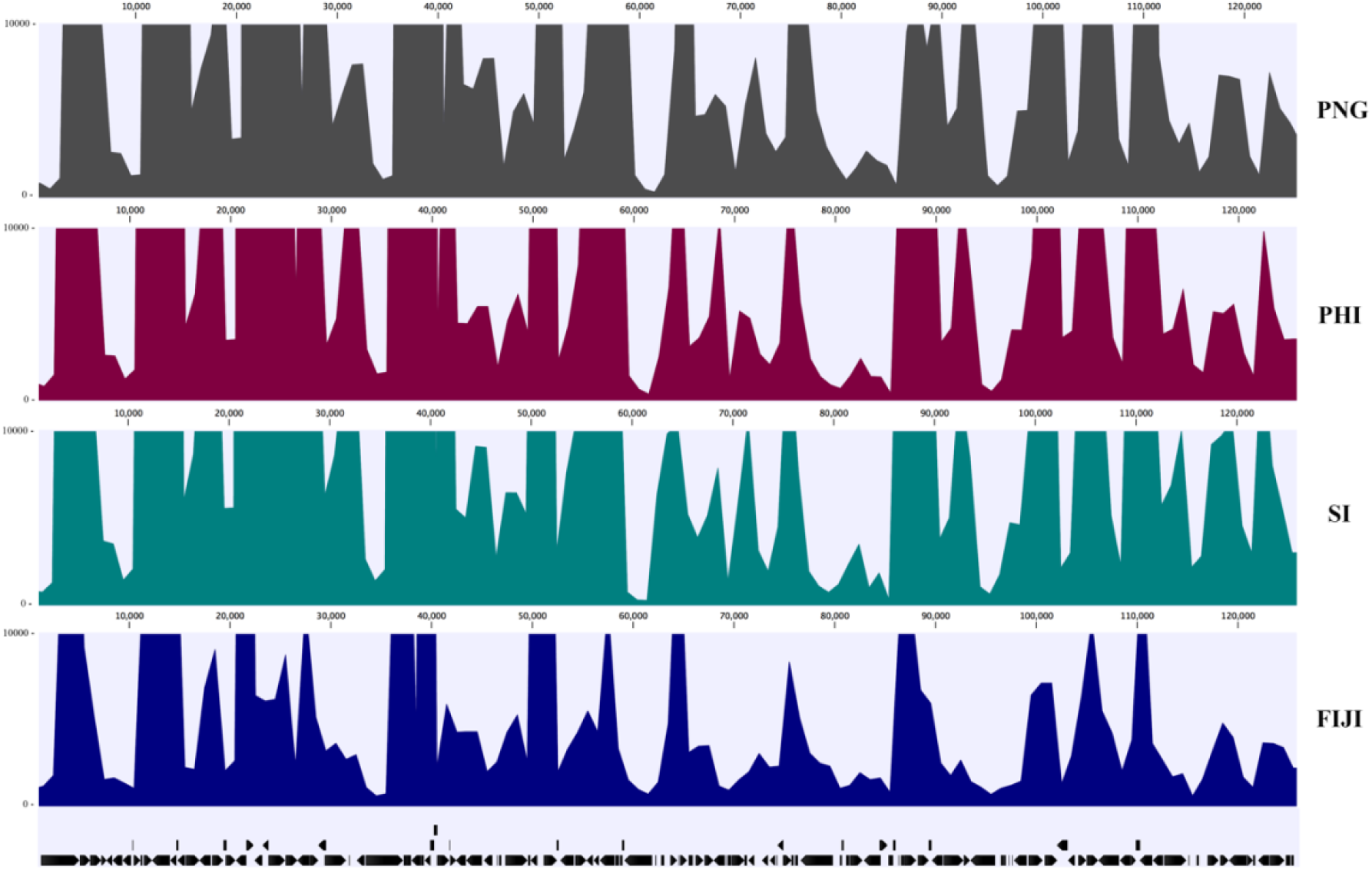
The average coverage of RNAseq data mapped to the OrNV genome (Solomon Islands Strain). A consensus sequence of virus was generated for each sample.

**Figure S2:**
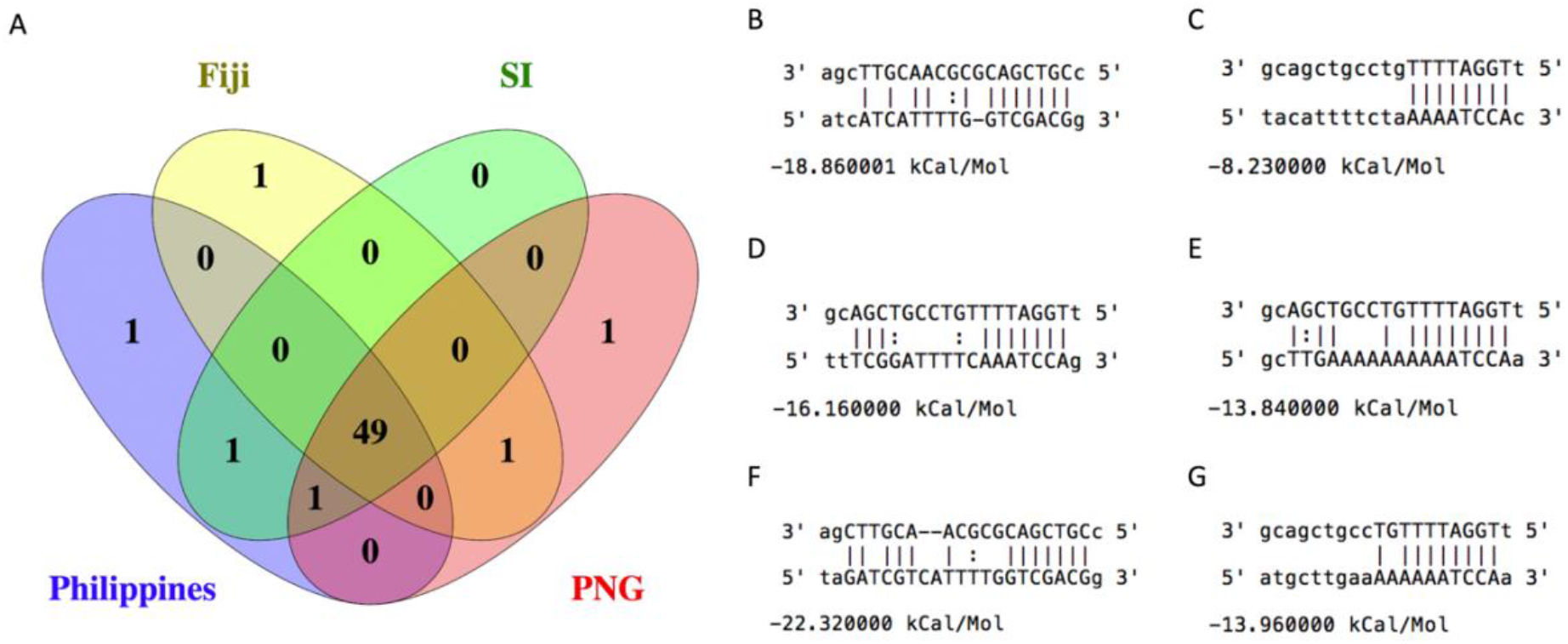
OrNv-miR-1 binding sites on the genome of different geographical strains of OrNV. Venn diagram represents the number of common and strain specific binding sites **(A)**. OrNv-miR-1-3p binding site on gp_027 exclusively in Fiji strains **(B)**; OrNv-miR-1-5p binding site on gp_069 exclusively in PNG strains **(C)**; OrNv-miR-1-5p binding site on gp_083 exclusively in Philippines strains **(D)**; A common OrNv-miR-1-5p binding site on a non-coding part of genome in Fiji and PNG strains **(E)**; A common OrNv-miR-1-3p binding site on gp_027 in Philippines, Solomon Islands and PNG strains **(F)**; A common OrNv-miR-1-5p binding site on a non-coding part of genome in Philippines and Solomon Islands strains **(G)**

